# Human APOBEC3G prevents emergence of infectious endogenous retrovirus in mice

**DOI:** 10.1101/457937

**Authors:** Rebecca S. Treger, Maria Tokuyama, Huiping Dong, Susan R. Ross, Yong Kong, Akiko Iwasaki

## Abstract

Endogenous retroviruses (ERV) are found throughout vertebrate genomes and failure to silence their activation can have deleterious consequences on the host. Introduction of mutations that subsequently prevent transcription of ERV loci is therefore an indispensable cell-intrinsic defense mechanism that maintains the integrity of the host genome. Abundant *in vitro* and *in silico* evidence have revealed that APOBEC3 cytidine-deaminases, including human APOBEC3G (hA3G) can potently restrict retrotransposition; yet *in vivo* data demonstrating such activity is lacking, particularly since no replication competent human ERV has been identified. In mice deficient for Toll-like receptor 7 (TLR7), transcribed ERV loci can recombine and generate infectious ERV. In this study, we show that mice deficient in the only copy of *Apobec3* in the genome did not have spontaneous reactivation of ERVs, nor elevated ERV reactivation when crossed to *Tlr7*^-/-^ mice. In contrast, expression of a human APOBEC3G transgene abrogated emergence of infectious ERV in the *Tlr7*^-/-^ background. No ERV RNA was detected in the plasma of hA3G+*Apobec3*-/-*Tlr7*^-/-^ mice, and infectious ERV virions could not be amplified through co-culture with permissive cells. These data reveal that hA3G can potently restrict active ERV *in* vivo, and suggest that the expansion of the APOBEC3 locus in primates has helped restrict ERV reactivation in the human genome.

**Importance:** Although APOBEC3 proteins are known to be important antiviral restriction factors in both mice and humans, their roles in the restriction of endogenous retroviruses (ERV) have been limited to *in vitro* studies. Here, we report that human APOBEC3G expressed as a transgene in mice prevents the emergence of infectious ERV from endogenous loci. This study reveals that APOBEC3G can powerfully restrict active retrotransposons *in vivo* and demonstrates how ectopic expression of human factors in transgenic mouse models can be used to investigate host mechanisms that inhibit retrotransposons and reinforce genomic integrity.

## Body

Roughly eight to ten percent of both human and murine genomes is composed of endogenous retroviruses (ERV), the endogenized counterparts of ancient retroviruses that invaded the germline and became fixed within these genomes (1, 2). The proviral-like ERV present in the genomes of common laboratory mouse strains formed following infection by an exogenous murine leukemia virus (MLV), and these ERV loci are actively transcribed and translated. Although wild-type C57BL/6 mice do not contain a proviral ERV locus capable of independently generating replication-competent ERV (3), infectious ERV virions readily emerge when B-cell elicited humoral control mediated through toll-like receptor 7 (TLR7) signaling is deficient (4, 5). As with emerged ERV from recombination-activating gene 1-deficient (*Rag1*^-/-^) mice (4), we observed that infectious ERV in *Tlr7*^-/-^ mice result from recombination between *Emv2*, the only ecotropic ERV in the C57BL/6 genome, and several endogenous retroviral loci (Figure 1A). Thus, as with proviruses, cell-intrinsic control of ERV is essential to limit their transcription and potential for recombination and emergence. In addition to stochastic recombination events that remove ERV from the genome and transcriptional silencing of ERV loci (6), mutagenesis of retroelement sequences by Apolipoprotein B editing complex 3 (APOBEC3) proteins is an important component of this innate defense against ERV.

**Figure 1.**
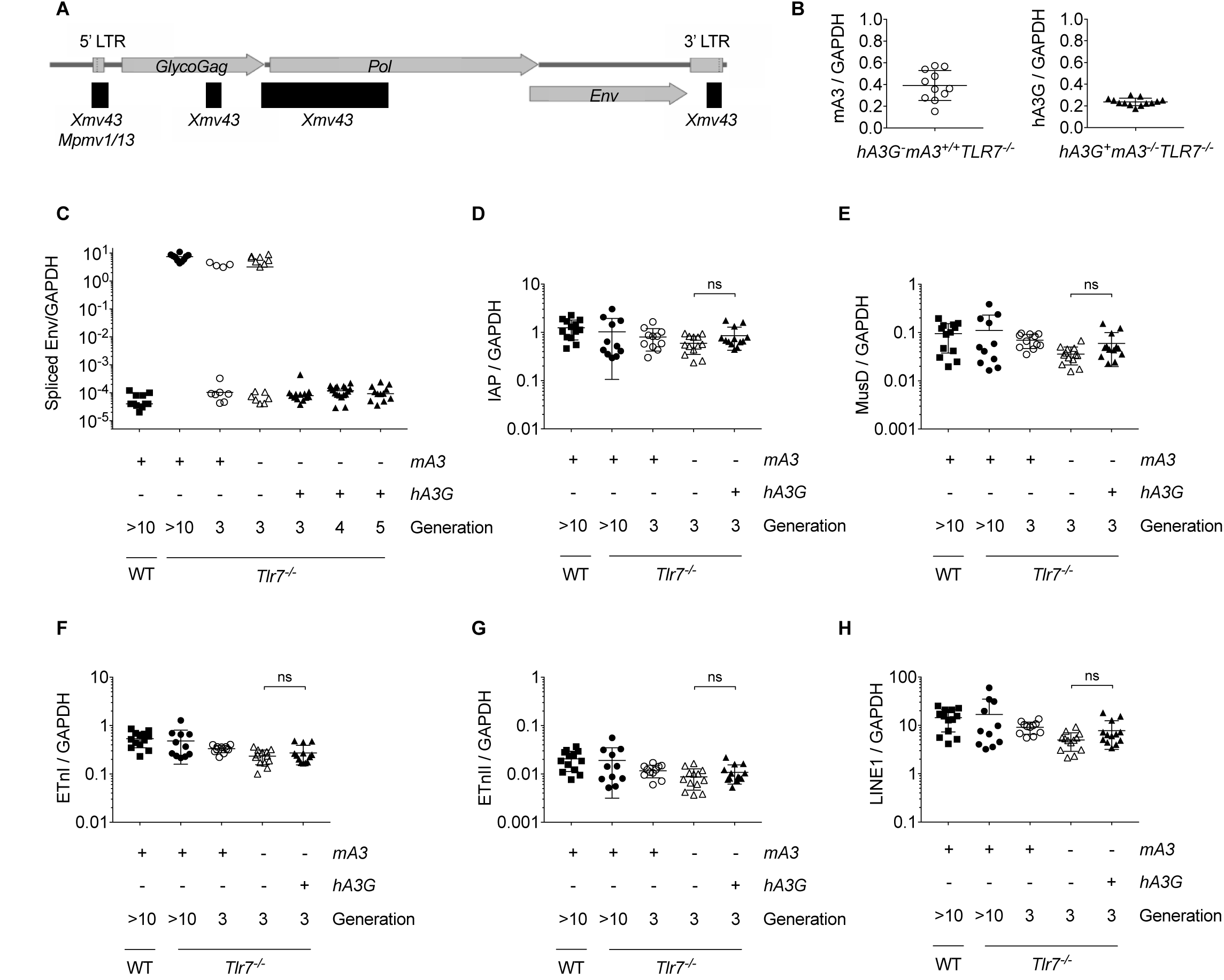
Human APOBEC3G, but not murine APOBEC3, expression prevents the emergence of infectious ERV in *Tlr7*^-/-^ mice. **A.** Schematic of the structure and open reading frames of the *Emv2*-based ERV genome isolated from virions amplified through co-culture with *Tlr7*^-/-^ splenocytes. Recombined regions are denoted by black horizontal bars and the ERV locus that contributed sequence is listed. **B.** RT-qPCR of *hA3G* and *mA3* expression from *hA3G+mA3-/-Tlr7^-/-^* (n=13) and *hA3G-mA3+/+Tlr7^-/-^* (n=11) mice, respectively. **C.** RT-qPCR of spliced *Emv2* envelope expression from C57BL/6N (n=13), *Tlr7*^-/-^ (n=11), F3 *hA3G-mA3+/+Tlr7^-/-^* (n=11), F3 *hA3G-mA3^-/-^Tlr7^-/-^* (n=15), F3 *hA3G+mA3-/-Tlr7^-/-^* (n=13), F4 *hA3G+mA3^-/-^Tlr7^-/-^* (n=18), and F5 *hA3G+mA3^-/-^Tlr7^-/-^* (n=12) mice. **D-H.** Expression of select LTR retrotransposon families via RT-qPCR using RNA isolated from peripheral blood of C57BL/6N (n=13), *Tlr7*^-/-^ (n=11), F3 hA3G-mA3+/+*Tlr7*^-/-^ (n=11), F3 hA3G-mA3-/-412 *Tlr7*^-/-^ (n=13), and F3 hA3G+mA3^-/-^*Tlr7*^-/-^ (n=13) mice. Primers amplify the gag or polymerase regions of IAP, MusD, and ETn elements (31), or LINE1 ORFp1. All values are normalized to GAPDH expression. Mean and standard deviation are plotted.

Present throughout vertebrate genomes, APOBEC3 proteins are zinc-dependent cytosine deaminases that act on single-stranded DNA to cause cytosine-to-uracil mutations in target sequences (7). Although mouse genomes encode a single *Apobec3* gene, *mA3*, expansion of this locus in primates has given rise to seven *APOBEC3* genes, *APOBEC3A*, -*3B*, -*3D/E*, -*3F*, -*3G*, & -*3H* (8). APOBEC3 proteins, and particularly the human APOBEC3G (hA3G), have long been appreciated for their potent restriction of exogenous retroviruses. Originally characterized for its activity against human immunodeficiency virus 1 (HIV-1) (9), hA3G is a restriction factor that is packaged into retroviral virions, which upon release into target cell hypermutates reverse transcribed viral ssDNA through its deaminase domain (10, 11). hA3G also inhibits reverse transcriptase in a deaminase-independent manner (12-14).

Additionally, hA3G restricts murine ERV when overexpressed in *in vitro* reporter assays (15-17), and can hypermutate human ERV (HERV) sequences (18, 19). This *in vitro* evidence is also supported by *in silico* data that mA3 and hA3 family members have targeted murine ERV and HERV genomic loci (20), respectively, including those encoding the proviral ERV capable of emergence in mice (21). Meanwhile, *in vivo* studies have demonstrated that mA3 restricts MLV (*22, 23*). In addition, hA3G is capable of blocking primary infection with exogenous MLV when expressed as a transgene in mice (24). Yet it remains unclear the extent to which hA3G restricts ERV and other retroelements *in vivo*, particularly since replication-competent HERV have not been identified in the human genome (25) and identification of A3-restricted retrotransposons is complicated by the high copy number and repetitive nature of the retroelements themselves. On the other hand, in C57BL/6 mice, a single proviral ERV, *Emv2*, forms the backbone of the infectious ERV that emerge (4). Because this locus is unique, its increased expression serves as an indicator of ERV emergence. We therefore took advantage of this phenomenon and available *hA3G* transgenic mice on the *mA3* knockout (*mA3*^-/-^) background (24) to investigate if mA3 and hA3G proteins are able to prevent or impede the emergence of replication competent ERV in *Tlr7*^-/-^ mice *in vivo*.

To investigate the role of mA3 and hA3G in the restriction of ERV, we first crossed *hA3G+* mice lacking *mA3* (*hA3G+mA3*^-/-^) to *Tlr7*^-/-^ mice to generate a first generation (F1) of transgene-positive and ‐negative mice with homozygous loss of TLR7 (*hA3G+mA3-/-*101 *Tlr7*^-/-^ and *hA3G-/mA3^-/-^Tlr7^-/-^*). We also bred mice that maintained mA3 expression in the absence of TLR7 (*hA3G-mA3+/+Tlr7^-/-^*). These three strains were then bred out for several generations and screened for both A3 expression (Figure 1B) and ERV emergence (Fig. 1C) by reverse transcription quantitative polymerase chain reaction (RT-qPCR).

Generation of replication-competent ERV requires multiple recombination events to restore polymerase function and endow the *Emv2*-based virus with a non-restricted capsid (3). Accordingly, we first observed ERV emergence in *mA3+/+Tlr7^-/-^* and *mA3*^-/-^*Tlr7*^-/-^ mice by the third generation (F3) of breeding between homozygous knockouts (Figure 1C). More than half (8/15) of the *mA3^-/-^Tlr7^-/-^* mice were ERV-positive, as were 4 of the 11 *mA3+/+Tlr7^-/-^* controls. These data indicated that mA3 is not sufficient to prevent emergence of ERV. Like most MLV, ERV express glycosylated Gag (Figure 1A), a longer, glycosylated variant of Gag protein that counters restriction by mA3 (22). This antagonization may underlie the failure of mA3 to prevent ERV emergence.

In stark comparison to the effect of mA3, hA3G expression in the *Tlr7*^-/-^ background entirely abrogated ERV emergence (Figure 1C). In the F3 offspring, all 13 *hA3G+mA3^-/-^Tlr7*^-/-^ mice retained control of ERV, and this impressive capacity to prevent ERV emergence extended to the fourth (F4) and to fifth (F5) generations, where all *hA3G+mA3^-/-^Tlr7^-/-^* mice remained infectious ERV-negative. We did not observe differences in global expression levels of IAP, MusD, early transposon I & II (ETnI & ETnII) elements, or LINE1 between any of the genotypes by RT-qPCR (Figure 1D-H). Thus, these data suggest that hA3G is highly specific for *Emv2*-derived ERV and is not involved in the widespread suppression of LTR retroelement or LINE1 transcription.

Restriction of MLV by hA3G involves both cytosine deamination-dependent and ‐independent mechanisms (24). We hypothesize that restriction of ERV by hA3G includes hypermutation of partially or fully recombinant ERV transcripts. We were able to amplify infectious ERV from *hA3G-Tlr7^-/-^* splenocytes in co-culture with permissive cell lines (Figure 2A-C) and to detect circulating ERV RNA in plasma from these mice (Figure 2D). However, we were unable to detect ERV RNA in the plasma of and were unable to amplify infectious ERV from *hA3G+mA3^-/-^Tlr7^-/-^* mice (Figure 2A-D). Because infectious ERV did not emerge in these mice, we are unable to ascertain the extent to which deamination contributes to restriction in transgene-positive mice. It is also possible that, as with MLV and HIV-1 infection, hA3G inhibits the reverse transcriptase of polymerase-restored ERV or otherwise impairs subsequent integration (26, 27) during the initial stages of emergence, before infectious virions are reconstituted. To study how hA3G specifically restricts ERV emergence would require the development of a new *in vitro* system that accurately recapitulates the events that give rise to emergence.

**Figure 2.**
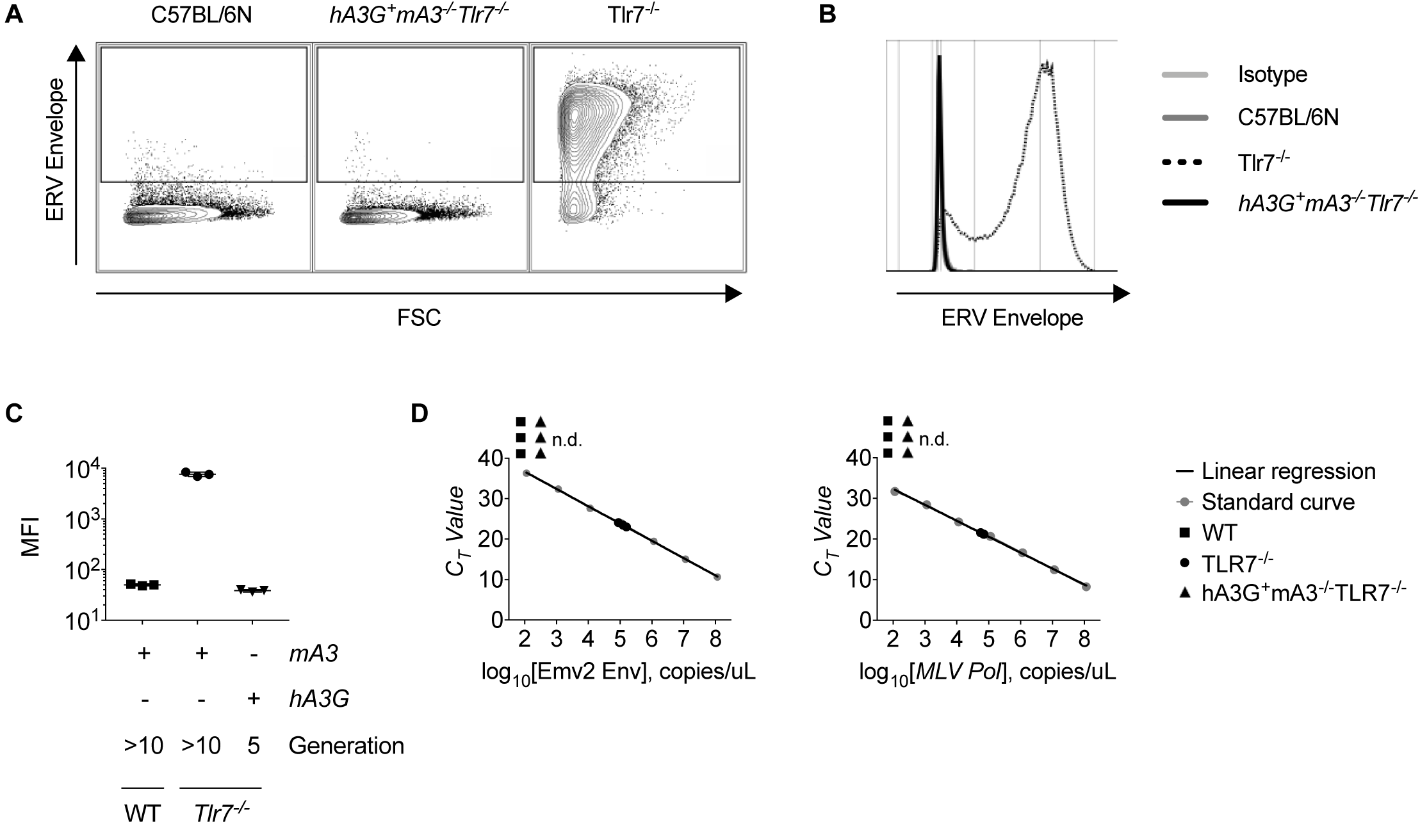
Infectious ERV cannot be detected in the plasma or isolated through splenocyte co-culture from hA3G+mA3-/-*Tlr7*^-/-^ mice. **A-C.** Representative flow cytometry plots (A), histograms (B), and calculated mean fluorescent intensities (C) of ERV envelope expression on live, CD45.2-negative DFJ8 cells following 7 days of co-culture with C57BL/6N, *Tlr7*^-/-^, or F5 *hA3G+mA3-/-Tlr7*^-/-^ splenocytes (n=3 mice per group). **D.** Absolute quantification of the number of polymerase or unspliced ERV envelope RNA copies per microliter of cDNA generated from plasma of 16-week-old C57BL/6N, *Tlr7*^-/-^, and *hA3G+mA3^-/-^Tlr7^-/-^* mice. Plots are representative of 3 independent experiments.

In this study, we have demonstrated that unlike mA3, hA3G powerfully restricts ERV *in vivo*, preventing the emergence of infectious ERV that would otherwise occur in the absence of TLR7 signaling. These data extend our understanding of the function of this protein and reveal an important layer of host defense that reinforces genomic integrity. Indeed, the expansion of the APOBEC3 locus and the presence of hA3G may contribute to the mechanisms that prevent reconstitution of replication-competent HERV in humans. While the sequence of events that results in ERV emergence has yet to be characterized, this study demonstrates that mice transgenic for human restriction factors can serve as a powerful tool to investigate how such proteins, including hA3G, interact with retroelements and restrict their movement within the genome.

## Methods

### Mice

C57BL/6N mice were obtained from Charles Rivers and bred in-house. *Tlr7*^-/-^ (C57BL/6N) mice (28) were bred-in house. *hA3G+mA3^-/-^* and *hA3G-mA3-/-* mice (24) were maintained by breeding between transgene-positive and/or -negative *mA3* knockout mice. All mice were housed in SPF conditions and care was provided in accordance with Yale University IACUC guidelines (protocol #10365). The *hA3G-mA3^-/-^Tlr7*^-/-^ mice were maintained as a separate line from the transgene-negative littermates of *hA3G+mA3^-/-^Tlr7^-/-^* crosses to ensure that the ERV transcripts and genomic loci in the *hA3G-mA3^-/-^Tlr7^-/-^* genome were not subject to effects of hA3G expression.

### Peripheral blood isolation

Mice were anesthetized and blood was obtained via retro-orbital bleed. Blood was collected with heparinized Natelson tubes (Fisher Scientific) into 8mM EDTA in PBS. For cellular RNA isolation, red blood cells were lysed with ACK lysis buffer (150mM NH_4_Cl, 1 M KHCO_3_, 0.1mM EDTA, pH 7.4) and cells were washed twice with PBS before addition of RLT buffer (Qiagen). Samples were stored at -80 prior to RNA isolation.

### Reverse transcription-quantitative polymerase chain reaction (RT-qPCR)

RNA was isolated from peripheral blood using the RNeasy Kit (Qiagen) and cDNA was synthesized using iScript™ cDNA Synthesis Kit (Bio-Rad). Quantitative PCR was performed using iTaq™ Universal SYBR^®^ Green Supermix (Bio-Rad) in 10ul reactions in triplicate using 5-30ng of cDNA per reaction. Primers were used at a final concentration of 0.225μM. Primer sequences are as follows:

Spliced Emv2 Env (4) Forward 5’-CCAGGGACCACCGACCCACCGT-3’ Reverse 5′-TAGTCGGTCCCGGTAGGCCTCG-3′

Unspliced Emv2 Env (29) Forward 5’-AGGCTGTTCCAGAGATTGTG-3’ Reverse 5’-TTCTGGACCACCACATGAC-3’

GAPDH Forward 5’-GAAGGTCGGTGTGAACGGA-3’ Reverse 5’-GTTAGTGGGGTCTCGCTCCT-3’

IAP (30) Forward 5’-AAGCAGCAATCACCCACTTTGG-3’ Reverse 5’-CAATCATTAGATGTGGCTGCCAAG-3’

MusD (30) Forward 5’-GTGGTATCTCAGGAGGAGTGCC-3’ Reverse 5’-GGGCAGCTCCTCTATCTGAGTG-3’

ETnI (31) Forward 5’-TGAGAAACGGCAAAGGATTTTTGGA-3’ Reverse 5’-ATTACCCAGCTCCTCACTGCTGA-3’

ETnII (31) Forward 5’-GTGCTAACCCAACGCTGGTTC-3’ Reverse 5’-ACTGGGGCAATCCGCCTATTC-3’

### Plasma RNA isolation and cDNA synthesis

Peripheral blood was isolated from 16-week-old mice and centrifuged at 14,000rpm for 15min at 4 degrees Celsius and 200uL of plasma was removed to a new Eppendorf tube. Plasma was homogenized with 1mL of Trizol and 200uL of chloroform, and the aqueous layer was isolated by centrifugation for 15min at 12,000*g* at 4 degrees Celsius. The aqueous layer was combined with 500uL isopropanol and 90ug/mL glycerol and frozen for 1hr at -80 degrees Celsius. The RNA was then pelleted by centrifugation for 10 min at 12,000*g* at 4 degrees Celsius and washed twice with cold 75% ethanol before resuspending in 10uL of RNase-free water. cDNA was synthesized using the Superscript III Cells Direct cDNA Synthesis Kit (Invitrogen) and qPCR was performed as above, using sequenced ERV plasmid (described below) to generate a standard curve for absolute quantification.

### Splenocyte Isolation and co-culture

The day before co-culture, 100,000 DFJ8 avian fibroblasts were plated in 1mL of DMEM (Gibco) supplemented with 10% FBS and 1% penicillin/streptomycin (Gibco) in a 12-well tissue culture-treated dish. On the day of co-culture, spleens were isolated and dissociated through a 40μm filter in RPMI media (Gibco) and red blood cells were lysed with ACK lysis buffer. The splenocytes were then washed and passed through a 70μm filter prior to counting, and 5 million splenocytes from each mouse were added to a corresponding well of DFJ8 cells, supplemented with an additional 1mL of media. Four days later, each co-culture (supernatant plus cells) was moved to a 60mm dish using 2mM EDTA in PBS to dissociate the adherent cells, with a final culture volume of 4mL. On day 7 of co-culture, the adherent cells were stained for ERV envelope expression by flow cytometry.

### Flow cytometry

Hybridoma supernatant containing monoclonal antibody 573 was a kind gift of Leonard Evans (32). Cells were washed twice with PBS and stained with mAb573 diluted 1:1 with PBS, then washed twice again and stained with anti-mouse CD45.2-FITC (BioLegend 109805), anti-mouse IgM-APC (Jackson 115-136-075), and 7-AAD viability staining solution (eBioscience 00-6993-50). Prior to analysis, cells were fixed in 1% paraformaldehyde in PBS. All incubations were performed at a final volume of 30μL for 15-20min at 4 degrees Celsius. Flow cytometry was performed on a BD LSRII Green cytometer and data was analyzed with FlowJo.

### ERV isolation and sequencing

Individual ERV-infected DFJ8 cells from co-culture with TLR7^-/-^ splenocytes were seeded in a 96-well plate and expanded until confluent in 12-well dishes. These monoclonal cultures were analyzed for ERV envelope expression by flow cytometry (as described above), and a single infected clone (D61) was selected. Two million D61 cells were plated in T-175 flasks in 30mL of media and grown for 1 week, after which the supernatant was harvested. Cell debris was removed by centrifuging the sample at 1500rpm for 5 min, and the resulting supernatant was clarified through a 0.45um filter. The clarified supernatant was underlaid with a 1.12 g/ul sucrose cushion and ultracentrifuged at 23,000rpm for 2hr at 4 degrees Celsius. The resulting viral pellet was resuspended in Opti-MEM (Gibco) and stored at -20 degrees Celsius. ERV RNA was isolated from these viral stocks using Trizol/Chloroform extraction and random hexamer cDNA synthesis (described above). Using the *Emv2* mm10 genomic sequence, primers to highly conserved regions of the *Emv2* backbone were used to amplify overlapping segments of the viral genome, which was assembled using Gibson Assembly (NEB) and cloned into the pUC19 vector for sequencing.

### ERV recombination analysis

A moving, overlapping window of size 50bp was used to extract fragment sequences from the *Emv2* ERV sequence. The step of the moving window is 1bp. The fragment sequences were used as queries to search the GRCm38 reference genome by blat with -fastMap option. Overused tile file with tile size 11 was used in the search. For each query sequence, all its hits were sorted based on score = (% identity * alignment length), chromosomes and positions; only those hits that have maximum score (including ties) were kept. Each hit on this hit list was searched in both directions to expand the hit to maximum length, and the chromosome position with the maximum length was used as the mapped position of the *Emv2* ERV sequence in the GRCm38 genome. Regions of the ERV sequence that mapped to non-*Emv2* hits and were greater than 10bp were considered to have recombined with Emv2. Any recombined positions corresponding to unique ERV xenotropic (Xmv), polytropic (Pmv), or modified polytropic (Mpmv) loci (21) were identified.

## Acknowledgements

We would like to thank Huiping Dong for maintaining the mouse strains used in this study. We also thank Dr. Leonard Evans for sharing the 573 hybridoma supernatant.

## Author contributions

R.T., M.T., and A.I designed the experiments; R.T. performed the experiments; Y.K. analyzed ERV sequence data; R.T., S.R., and A.I. analyzed data; R.T. and A.I. prepared the manuscript.

## References

1. Lander ES, Linton LM, Birren B, Nusbaum C, Zody MC, Baldwin J, Devon K, Dewar K, Doyle M, FitzHugh W, Funke R, Gage D, Harris K, Heaford A, Howland J, Kann L, Lehoczky J, LeVine R, McEwan P, McKernan K, Meldrim J, Mesirov JP, Miranda C, Morris W, Naylor J, Raymond C, Rosetti M, Santos R, Sheridan A, Sougnez C, Stange-Thomann Y, Stojanovic N, Subramanian A, Wyman D, Rogers J, Sulston J, Ainscough R, Beck S, Bentley D, Burton J, Clee C, Carter N, Coulson A, Deadman R, Deloukas P, Dunham A, Dunham I, Durbin R, French L, Grafham D, et al. 2001. Initial sequencing and analysis of the human genome. Nature 409:860–921.

2. Waterston RH, Lindblad-Toh K, Birney E, Rogers J, Abril JF, Agarwal P, Agarwala R, Ainscough R, Alexandersson M, An P, Antonarakis SE, Attwood J, Baertsch R, Bailey J, Barlow K, Beck S, Berry E, Birren B, Bloom T, Bork P, Botcherby M, Bray N, Brent MR, Brown DG, Brown SD, Bult C, Burton J, Butler J, Campbell RD, Carninci P, Cawley S, Chiaromonte F, Chinwalla AT, Church DM, Clamp M, Clee C, Collins FS, Cook LL, Copley RR, Coulson A, Couronne O, Cuff J, Curwen V, Cutts T, Daly M, David R, Davies J, Delehaunty KD, Deri J, Dermitzakis ET, et al. 2002. Initial sequencing and comparative analysis of the mouse genome. Nature 420:520–562.

3. Young GR, Kassiotis G, Stoye JP. 2012. Emv2, the only endogenous ecotropic murine leukemia virus of C57BL/6J mice. Retrovirology 9:23.

4. Young GR, Eksmond U, Salcedo R, Alexopoulou L, Stoye JP, Kassiotis G. 2012. Resurrection of endogenous retroviruses in antibody-deficient mice. Nature 491:774–778.

5. Yu P, Lubben W, Slomka H, Gebler J, Konert M, Cai C, Neubrandt L, Prazeres da Costa O, Paul S, Dehnert S, Dohne K, Thanisch M, Storsberg S, Wiegand L, Kaufmann A, Nain M, Quintanilla-Martinez L, Bettio S, Schnierle B, Kolesnikova L, Becker S, Schnare M, Bauer S. 2012. Nucleic acid-sensing Toll-like receptors are essential for the control of endogenous retrovirus viremia and ERV-induced tumors. Immunity 37:867–879.

6. Goodier JL. 2016. Restricting retrotransposons: a review. Mobile DNA 7:16.

7. Conticello SG. 2008. The AID/APOBEC family of nucleic acid mutators. Genome Biol 9:229.

8. Koito A, Ikeda T. 2012. Apolipoprotein B mRNA-editing, catalytic polypeptide cytidine deaminases and retroviral restriction. Wiley Interdiscip Rev RNA 3:529–541.

9. Sheehy AM, Gaddis NC, Choi JD, Malim MH. 2002. Isolation of a human gene that inhibits HIV-1 infection and is suppressed by the viral Vif protein. Nature 418:646–650.

10. Lecossier D, Bouchonnet F, Clavel F, Hance AJ. 2003. Hypermutation of HIV-1 DNA in the absence of the Vif protein. Science 300:11–12.

11. Zhang H, Yang B, Pomerantz RJ, Zhang C, Arunachalam SC, Gao L. 2003. The cytidine deaminase CEM15 induces hypermutation in newly synthesized HIV-1 DNA. Nature 424:94–98.

12. Guo F, Cen S, Niu M, Saadatmand J, Kleiman L. 2006. Inhibition of tRNA(3)(Lys)-primed reverse transcription by human APOBEC3G during human immunodeficiency virus type 1 replication. J Virol 80:11710–11722.

13. Guo F, Cen S, Niu M, Yang Y, Gorelick RJ, Kleiman L. 2007. The interaction of APOBEC3G with human immunodeficiency virus type 1 nucleocapsid inhibits tRNA3Lys annealing to viral RNA. J Virol 81:11322–11331.

14. Stavrou S, Ross SR. 2015. APOBEC3 Proteins in Viral Immunity. Journal of immunology (Baltimore, Md: 1950) 195:4565–4570.

15. Esnault C, Heidmann O, Delebecque F, Dewannieux M, Ribet D, Hance AJ, Heidmann T, Schwartz O. 2005. APOBEC3G cytidine deaminase inhibits retrotransposition of endogenous retroviruses. Nature 433:430–433.

16. Esnault C, Millet J, Schwartz O, Heidmann T. 2006. Dual inhibitory effects of APOBEC family proteins on retrotransposition of mammalian endogenous retroviruses. Nucleic Acids Research 34:1522–1531.

17. Smith HC, Bennett RP, Kizilyer A, McDougall WM, Prohaska KM. 2012. Functions and Regulation of the APOBEC Family of Proteins. Seminars in cell & developmental biology 23:258–268.

18. Lee YN, Malim MH, Bieniasz PD. 2008. Hypermutation of an Ancient Human Retrovirus by APOBEC3G. Journal of Virology 82:8762–8770.

19. Esnault C, Priet S, Ribet D, Heidmann O, Heidmann T. 2008. Restriction by APOBEC3 proteins of endogenous retroviruses with an extracellular life cycle: ex vivo effects and in vivo”traces” on the murine IAPE and human HERV-K elements. Retrovirology 5:75.

20. Anwar F, Davenport MP, Ebrahimi D. 2013. Footprint of APOBEC3 on the Genome of Human Retroelements. Journal of Virology 87:8195–8204.

21. Jern P, Stoye JP, Coffin J. 2005. Role of APOBEC3 in Genetic Diversity among Endogenous Murine Leukemia Viruses. PLoS Genetics doi:10.1371/journal.pgen.0030183.eor:e183.

22. Harris RS, Dudley JP. 2015. APOBECs and virus restriction. Virology 479-480:131–145.

23. Low A, Okeoma CM, Lovsin N, de las Heras M, Taylor TH, Peterlin BM, Ross SR, Fan H. 2009. Enhanced replication and pathogenesis of Moloney murine leukemia virus in mice defective in the murine APOBEC3 gene. Virology 385:455–463.

24. Stavrou S, Crawford D, Blouch K, Browne EP, Kohli RM, Ross SR. 2014. Different Modes of Retrovirus Restriction by Human APOBEC3A and APOBEC3G In Vivo. PLoS Pathogens 10:e1004145.

25. Kassiotis G. 2014. Endogenous Retroviruses and the Development of Cancer. The Journal of Immunology 192:1343.

26. Luo K, Wang T, Liu B, Tian C, Xiao Z, Kappes J, Yu X-F. 2007. Cytidine Deaminases APOBEC3G and APOBEC3F Interact with Human Immunodeficiency Virus Type 1 Integrase and Inhibit Proviral DNA Formation. Journal of Virology 81:7238–7248.

27. Mbisa JL, Barr R, Thomas JA, Vandegraaff N, Dorweiler IJ, Svarovskaia ES, Brown WL, Mansky LM, Gorelick RJ, Harris RS, Engelman A, Pathak VK. 2007. Human Immunodeficiency Virus Type 1 cDNAs Produced in the Presence of APOBEC3G Exhibit Defects in Plus-Strand DNA Transfer and Integration. Journal of Virology 81:7099–7110.

28. Lund JM, Alexopoulou L, Sato A, Karow M, Adams NC, Gale NW, Iwasaki A, Flavell RA. 2004. Recognition of single-stranded RNA viruses by Toll-like receptor 7. Proc Natl Acad Sci U S A 101:5598–5603.

29. Yoshinobu K, Baudino L, Santiago-Raber ML, Morito N, Dunand-Sauthier I, Morley BJ, Evans LH, Izui S. 2009. Selective up-regulation of intact, but not defective env RNAs of endogenous modified polytropic retrovirus by the Sgp3 locus of lupus-prone mice. J Immunol 182:8094–8103.

30. Matsui T, Leung D, Miyashita H, Maksakova IA, Miyachi H, Kimura H, Tachibana M, Lorincz MC, Shinkai Y. 2010. Proviral silencing in embryonic stem cells requires the histone methyltransferase ESET. Nature 464:927–931.

31. Maksakova IA, Zhang Y, Mager DL. 2009. Preferential epigenetic suppression of the autonomous MusD over the nonautonomous ETn mouse retrotransposons. Mol Cell Biol 29:2456–2468.

32. Evans LH, Boi S, Malik F, Wehrly K, Peterson KE, Chesebro B. 2014. Analysis of Two Monoclonal Antibodies Reactive with Envelope Proteins of Murine Retroviruses: One Pan Specific Antibody and One Specific for Moloney Leukemia Virus. Journal of virological methods 200:47–53.

